# The human Flower isoform hFWE4 facilitates cornification in cutaneous squamous cell carcinoma

**DOI:** 10.1101/2025.09.10.671985

**Authors:** Justin C. Rudd, Patrick T. Kuwong, Rachel E. Johnson, Louise N. Monga-Wells, Meghan Vo, Julia Russolillo, Shreya Reddy, Mallory Jacobs, Hunter Litz, Changzhao Li, Andrew Siref, James A. Grunkemeyer, Laura A. Hansen

**Affiliations:** Department of Biomedical Sciences, Creighton University School of Medicine, Omaha, Nebraska, USA; Department of Pathology, Creighton University School of Medicine, Omaha, Nebraska, USA

## Abstract

Cutaneous squamous cell carcinoma (cSCC) is the second most common human tumor and arises from keratinocytes that comprise the epidermis and its associated hair follicles. Remarkably, transformed keratinocytes within a cSCC retain some ability to execute the terminal differentiation program that typically generates a stratified epidermis. Instead of the polarized stratification towards the body surface seen in normal epidermis, cSCC keratinocytes differentiate concentrically inward to produce keratin pearls. It is well appreciated that cSCC differentiation status is correlated with degree of local invasion and potential for distant metastasis, with well differentiated tumors having a better prognosis. While molecular mechanisms that govern the differentiation process are incompletely understood, identifying candidate molecules associated with cSCC differentiation can provide new histologic markers for prognostic stratification. Here we demonstrate that Flower (FWE), a newly described regulator of lamellar body (LB) trafficking in normal epidermis, is specifically expressed within highly differentiated layers of cSCC. Genetic knockout of the *hFWE* gene dysregulates cornification and LB-related gene expression leading to abnormal keratinization patterns, while ectopic hFWE4 expression drives G1-arrest and exit from the proliferative basal keratinocyte population. Additionally, we find that poorly differentiated keratinocyte regions of human cSCC exhibit minimal FWE-positivity, while this population is readily detectable in well-differentiated regions. We propose that FWE represents a novel differentiation marker in cSCC that could be utilized for classifying differentiation status and prognostic stratifying these tumors.

## INTRODUCTION

Proper execution of the terminal differentiation program in epidermal keratinocytes enables the formation of a functional barrier at the outer body surface (1). Upon commitment to differentiation, progenitor keratinocytes exit the proliferative basal layer and transit apically, undergoing extensive structural and biochemical modification before being sloughed off as anucleate corneocytes at the skin surface (2).

Abnormalities in this process typify a wide range of epidermal pathologies including epidermal differentiation disorders (3, 4), psoriasis, and cutaneous squamous cell carcinoma (cSCC) (5). Arising from transformed keratinocytes in the epidermis or associated hair follicles (6), cSCC is the second most common human cancer with over 700,000 cases reported annually in the United States (7). These tumors often retain an architecture that mimics the histology of normal epidermis with a basal proliferative population supplying overlying differentiated suprabasal layers and “keratin pearls” that are produced by terminally differentiated keratinocytes. While metastatic spread is rare and surgical excision of these tumors is often curative, certain patient demographics and histopathologic features carry a less favorable prognosis. Among histopathologic features, cSCC differentiation status is readily appreciated as a prognostic marker (8). Well-differentiated (WD) cSCC are characterized by extensive keratinization and formation of orderly differentiated structures called keratin pearls. WD cSCC carry a favorable prognosis while poorly-differentiated (PD) cSCC exhibit significantly higher rates of distant metastasis and mortality (8). Despite its clinical importance, grading of differentiation status in cSCC is highly subjective and several recent reports highlight a large degree of inter-rater variability (9–11). Molecular diagnostics such as immunohistochemistry for differentiation markers that can reliably discriminate between WD and PD cSCC promise to add precision to this process. However, identification of new markers for prognostic stratification requires a more complete understanding of the terminal differentiation process in cSCC.

Recently, we described a small four transmembrane protein, Flower (FWE), as a previously unappreciated regulator of lamellar body (LB) trafficking during the final stages of epidermal differentiation (12). While FWE has now been described as a regulator of vesicular and lysosome-related organelle trafficking (12–17) in several non-cutaneous tissues and cell types, it is not known whether FWE contributes to terminal differentiation in cSCC. Here, we demonstrate that FWE is expressed in highly differentiated keratinocytes in cSCC, that FWE loss is associated with alteration in the typical keratinization pattern, and that ectopic FWE expression facilitates exit from the proliferative basal population. Immunofluorescence on human cSCC reveals that FWE may serve as a novel differentiation marker to improve grading of cSCC differentiation status for prognostic purposes.

## RESULTS

To determine whether FWE expression is associated with differentiation in cSCC, we differentiated TERT-immortalized human epidermal keratinocytes (N/TERT-2G) alongside three cSCC cell lines, SCC-12b.2 (18, 19), SCC-13 (18, 19) and COLO16 (20), with variable potential for differentiation. Robust Ca^2+^-dependent expression of differentiation markers has been reported in SCC-12b.2 (21) which have minimal tumor forming capacity (22), while SCC-13 cells which readily form tumors (23, 24) exhibit only modest increase in total keratin content (25). After five days in 1.4mM Ca^2+^ containing EpiLife media, immunoblotting revealed a large increase in levels of Keratin 10 (K10) in N/TERT-2 and SCC-12b.2, while only a modest K10 response was observed in COLO 16 and was absent in SCC-13 (Fig 1A-B). Conversely, FWE expression was induced in all cell lines, with the most robust responses being observed in N/TERT-2G and SCC-12b.2 (Fig 1A-B). To gain further insight into the distribution of FWE expression during differentiation in cSCCs, we performed FWE immunofluorescence on subcutaneous SCC-13 xenografts. Although in monolayer culture, SCC-13 cells did not respond to calcium-induced differentiation by expressing K10, xenografts formed orderly differentiated structures called keratin pearls (Fig 1C). Immunofluorescence revealed that FWE signal was largely restricted to suprabasal nucleated keratinocyte layers before pearl formation (Fig 1C), consistent with the localization to upper stratum spinosum and granular layer keratinocytes in normal epidermis. Collectively, these findings suggested that as in normal epidermis, FWE is expression is induced during differentiation of cSCC keratinocytes.

**Figure 1.**
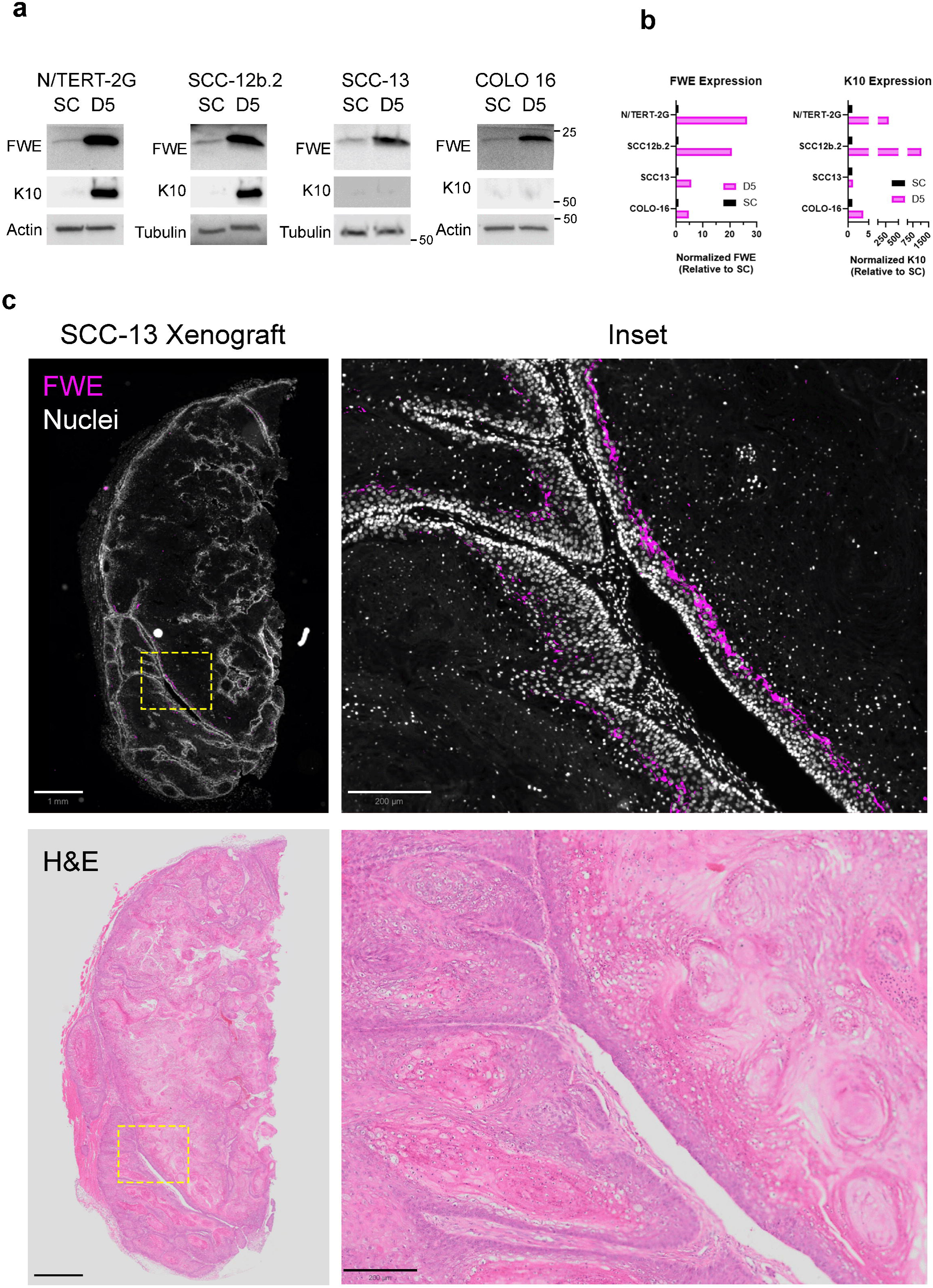
FWE is expressed by differentiated keratinocytes in cutaneous squamous cell carcinoma (cSCC). (A) Immunoblots of whole cell lysates from N/TERT-2G, SCC12b.2, SCC13 or Colo16 cells in the undifferentiated state or following 5d of differentiation in high calcium media. (B) Densitometric quantification of FWE and K10 immunoblots, presented as fold change relative to subconfluent. (C) Representative anti-FWE immunostaining in SCC13-derived xenograft (scale bar 1mm or 0.2mm (inset)).

We next questioned whether FWE expression contributed to proper execution of late-stage differentiation in cSCC. To test this question, we used CRISPR-Cas9 to knock out endogenous *hFWE* in SCC-13 cells, which have tumor forming capacity *in-vivo*, and isolated monoclonal cell lines (Fig 2A). Following validation of KO (Fig 2A, Supplementary Fig 1) control or *hFWE* KO SCC-13 cells were subcutaneously xenografted onto immunocompromised NCG mice and tumor volume was measured over 30 days. Xenografts derived from KO cell lines demonstrated a significant reduction in percent tumor area positive for FWE (Fig 2B). While no significant difference was observed in tumor growth (Fig 2C), a small but significant increase in mass of *hFWE* KO tumors was noted relative to WT controls (Fig 2D). This mass difference was accompanied by significant alterations in the keratinization pattern of *hFWE* deficient tumors relative to WT controls. Notably, keratin pearls in *hFWE* deficient tumors showed reduction in suprabasal cells marked by the presence of keratohyalin granules (Fig 2E, Supplementary Figure 2). *hFWE* deficient tumors also frequently exhibited solid parakeratosis at the center of keratin pearls, suggesting incomplete execution of the final stages of corneocyte formation (Fig 2E, Supplementary Figure 2). To validate these findings, immunofluorescence for late-stage differentiation markers filaggrin (Fig 2F) and loricrin (Fig 2G) was performed, revealing a significant reduction in total tumor area positive for these markers in *hFWE* KO tumors (Fig 2H). Conversely, knockout of *hFWE* had no effect on proliferation as measured by Ki67 positivity (Supplementary Figure 3). These findings suggested that as previously described in normal epidermis (12), *hFWE* expression is critical for execution of the latest stages of terminal differentiation in cSCC.

**Figure 2.**
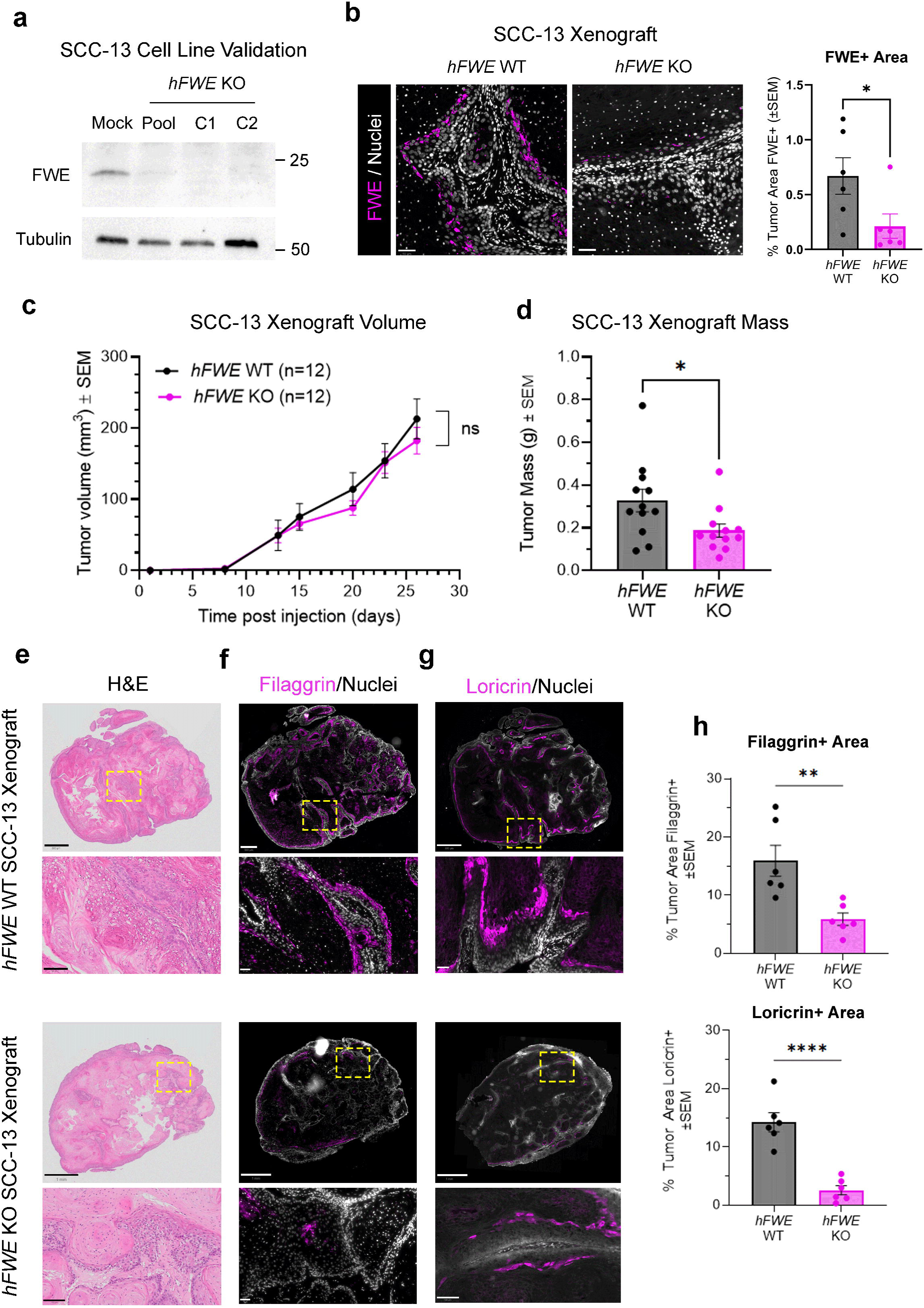
*FWE* KO alters keratinization pattern and impairs terminal differentiation in cSCC xenografts. (A) Representative FWE immunoblot of whole cell lysates from undifferentiated SCC13 cells. ‘Mock’ condition indicates electroporation with Cas9 alone, ‘Pool’ indicates polyclonal cell pool from Cas9:sgRNA electroporation, ‘C1’ and ‘C2’ indicate two distinct clonal cell lines derived from KO Pool. (B) Representative quantitative FWE immunofluorescence on xenografts derived from WT or KO SCC13 cells (scale bar 50 µm; two-tailed unpaired t-test, *p<0.05). (C) Tumor volumes of *hFWE* WT and KO SCC13 xenografts (n=12 biological replicates across three distinct clonal cell lines per genotype, multiple unpaired t-tests with post Benjamini, Krieger and Yekutieli correction). (D) Tumor mass of *hFWE* WT and KO SCC13 xenografts at necropsy (two-tailed unpaired t-test, *p < 0.05). (E) Representative H&E staining of *hFWE* WT (scale bar 800 µm, 100 µm (inset)) and KO xenografts (scale bar 1mm, 100 µm (inset)). (F) Representative filaggrin immunofluorescence of *hFWE* WT (scale bar 500 µm, 50 µm (inset)) and KO xenografts (scale bar 1mm, 50 µm (inset)). (G) Representative loricrin immunofluorescence of *hFWE* WT (scale bar 800 µm, 50 µm (inset)) and KO xenografts (scale bar 1mm, 100 µm (inset)). (H) Quantification of percent tumor area positive for filaggrin and loricrin in *hFWE* WT and KO xenografts (two-tailed unpaired t-test, **p<0.01, ****p<0.0001).

To gain additional insight into the molecular consequences of *hFWE* KO that might influence late terminal differentiation and cornification, we performed bulk RNA-seq of WT (N=12) and KO (N=11) tumors. Differential expression analysis revealed significant (FDR<0.05, l2fc>1) downregulation of 73 genes, while only four genes exhibited significant upregulation (Fig 3A, Supp Table 1). The significantly downregulated group was largely composed of genes with well described functions during the final stages of epidermal cornification that occur at the transition from stratum granulosum to stratum corneum. Notably, this included many kallikreins (*KLK5, KLK7, KLK8, KLK9, KLK10, KLK11, KLK12*), late-cornified envelope (LCE) family members (*LCE1B, LCE1C, LCE2A, LCE6A*), Secreted LY6/PLAUR Domain Containing proteins (*SLURP1, SLURP2*) and S100 family members (*S100A7, S100A12*). Gene ontology analysis of these 73 differentially downregulated genes demonstrated marked enrichment for biological processes related to keratinocyte differentiation and antimicrobial activity (Fig 3B), owing to the antimicrobial properties associated with many of these secreted molecules. Downregulated genes were also highly enriched for markers of the lamellar body (LB) compartment (Fig 3B) such as *KLK5* and *KLK7*, consistent with our previous work describing the role of FWE in LB biogenesis and secretion. Validating these RNA-seq findings, immunofluorescent signal for KLK5, a canonical cargo contained within FWE-positive LBs in normal human epidermis (12), was markedly reduced in differentiated regions of *hFWE* KO xenografts relative to WT controls (Fig 3C-D). Collectively, these data suggest loss of *hFWE* in cSCC dysregulates the final stages of cornification, likely by impairing the biogenesis and secretion of LBs.

**Figure 3.**
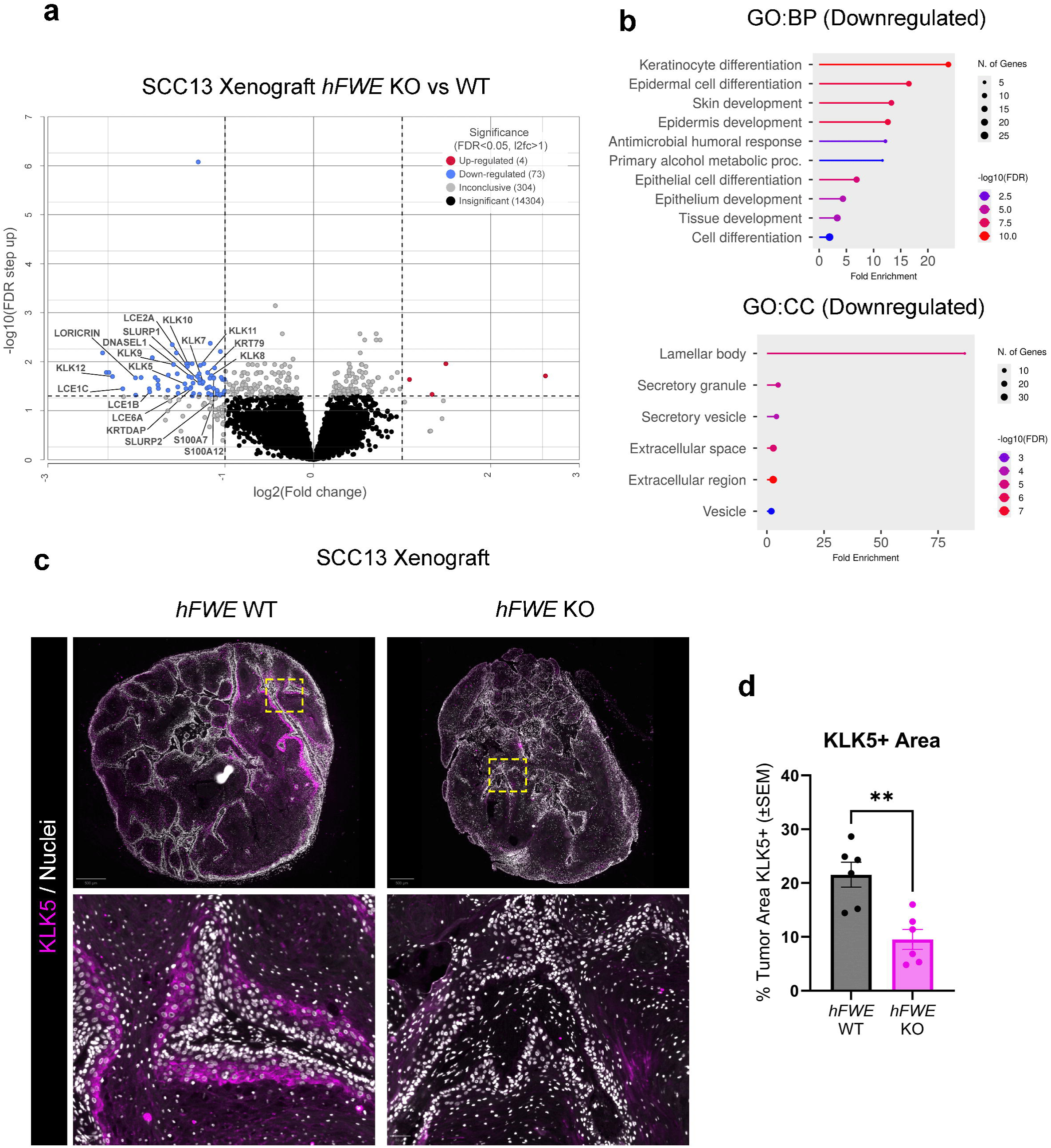
*FWE* KO dysregulates cornification and lamellar body related gene expression in cSCC xenografts. (A) Volcano plot representing differential expression analysis of bulk RNA-seq of *hFWE* WT (N=12) and KO (N=11) SCC13 xenografts. Genes implicated in lamellar body function or cornification are annotated. (B) GO:BP and GO:CC analysis on differentially downregulated genes in *hFWE* KO SCC13 xenografts. (C) Representative immunofluorescence for KLK5 in *hFWE* WT and KO SCC13 xenografts (scale bar 500 µm, 50 µm (inset)). (D) Quantification of percent tumor area KLK5 positive in *hFWE* WT and KO SCC13 xenografts (two-tailed unpaired t-test, **p<0.01).

Previously, we found that ectopic expression of the canonical hFWE isoform, hFWE4, in normal keratinocytes was sufficient to elicit cell cycle arrest in monolayer culture and exit from the proliferative basal compartment in organotypic epidermis (12). Given the impact of *hFWE* deficiency on keratinization in cSCCs, we reasoned that ectopic hFWE4 might similarly facilitate differentiation in these tumors. To test this hypothesis, we first introduced expression of EGFP alone or co-expressed EGFP with hFWE4-3XFLAG in a panel of cSCC cell lines (Fig 4A) and assayed cell cycle profiles of EGFP-positive cells in vitro. Across all cell lines, we found a significant increase in G1 and decrease in S-phase cells expressing hFWE4 compared to EGFP suggesting a G1-arrest (Fig 4B) similar to what we previously observed in normal keratinocytes. Next, we xenografted 1:1 mixtures of SCC13 cells expressing either EGFP alone or EGFP 2A hFWE4-3XFLAG with SCC13 cells expressing mCherry to assess how hFWE4 expression influenced the fate of keratinocytes in cSCCs (Fig 4C). No significant differences in tumor growth or mass were observed between groups (Supplementary Fig 4), Intriguingly, despite this lack of difference in tumor growth, immunostaining revealed that hFWE4 expressing cells were largely outcompeted by WT-mCherry cells (Fig 4E). Accordingly, immunofluorescence revealed that hFWE4-positive cells were significantly underrepresented in the proliferative (Ki67+) and basal (ITGB1+) populations, while they were significantly overrepresented in the highly differentiated (Filaggrin+) but not early differentiated (K10+) population (Fig 4F). Together, these findings indicate that much like in normal epidermis, ectopic expression of the canonical hFWE4 isoform is sufficient to drive G1-arrest and terminal differentiation of basal keratinocytes in cSCC.

**Figure 4.**
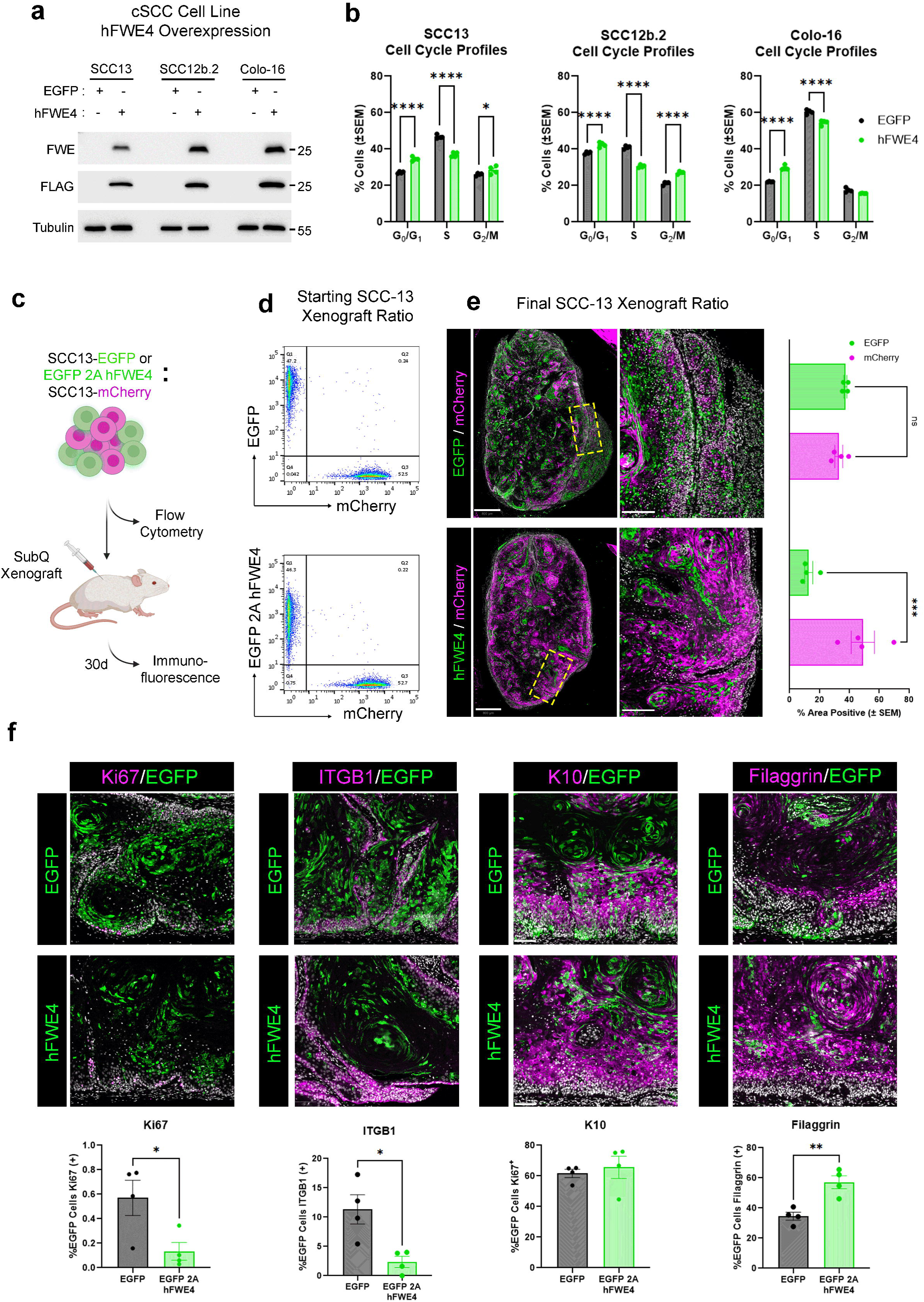
Ectopic hFWE4 expression facilitates cell cycle arrest and terminal differentiation in cSCC. (A) Immunoblot validation of EGFP 2A hFWE4-3XFLAG overexpression in SCC13, SCC12b.2 and Colo-16 cells. (B) Quantification of EGFP+ cells in indicated cell cycle stages following lentiviral infection with either EGFP or EGFP 2A hFWE4-3XFLAG (hFWE4) (n=4 biological replicates, *p<0.05, ****p<0.0001 2way ANOVA with Sidak’s multiple comparison test). (C) Schematic of experimental workflow for subcutaneous xenografting of 1:1 cocultures. (D) Flow cytometric analysis of EGFP+:mCherry+ ratio in cell suspensions used for xenografting. (E) Representative immunofluorescence for EGFP and mCherry in coculture xenografts grown for 30d in NCG mice. Quantification of percent EGFP or mCherry positive area across different coculture xenograft conditions (n=4, *p<0.05, **p<0.01 2way ANOVA with Sidak’s multiple comparison testing). (F) Representative immunofluorescence for EGFP and Ki67, ITGB1, K10 or Filaggrin in coculture xenografts. Quantification of the fraction of EGFP cells positive for the indicated marker across different xenograft conditions (*p<0.05, **p<0.01 two-tailed, independent t-test).

Based on the apparent association between hFWE and terminal differentiation in our cSCC models, we next sought to determine whether the presence of hFWE could serve as a reliable identifier of human cSCC differentiation status. To test this hypothesis, we performed FWE immunofluorescence on human cSCC identified as well-differentiated (n=9) or poorly differentiated (n=5) by a pathologist at the time of diagnosis. Following slide digitization, coverslips were removed, and sections were H&E stained in order to register FWE signal with tumor histology. Using H&E stains, regions of well-, moderately-, and poorly-differentiated keratinocytes within these tumors were subsequently identified and annotated by a dermatopathologist who was blinded to the corresponding FWE immunofluorescence. Regions identified as adjacent normal skin or actinic keratosis (AK) were annotated as well. Interestingly, we found that despite an initial designation as ‘well’ or ‘poorly-differentiated’ the degree of keratinocyte differentiation on subsequent annotation often varied within each tumor (Fig 5A), suggesting that differentiation grading might be refined through the use of molecular markers. Quantification of thresholded signal from FWE immunofluorescence showed that both poorly- and moderately-differentiated keratinocyte regions had significantly less FWE-positive area than well-differentiated regions, adjacent normal skin or AK (Fig 5B-C). Similarly, mean fluorescence intensity for FWE was significantly reduced in poorly- and moderately-differentiated keratinocyte regions (Fig 5B-C). Collectively, these data suggest that FWE is associated with terminal differentiation in human cSCC and that signal intensity may be a beneficial tool for accurate stratification of cSCC based on differentiation status for prognostic purposes.

**Figure 5.**
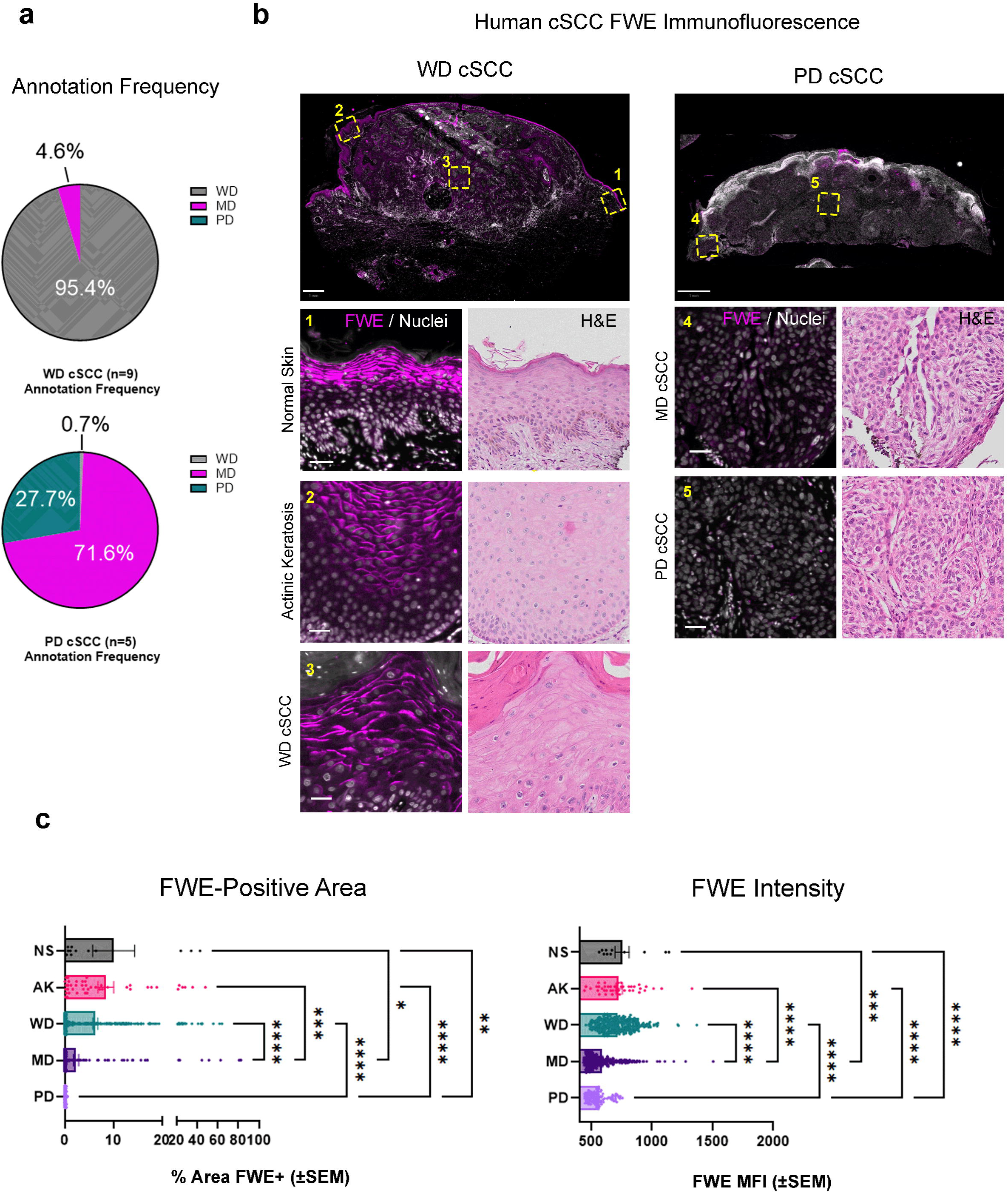
FWE expression is associated with differentiation status of human cSCC. (A) Separate pie charts representing patient cSCC biopsies initially classified as well-differentiated (WD, (N=9) (top)) or poorly-differentiated (PD, (N=5) (bottom)) at time of diagnosis. Colors indicate keratinocyte regions within these tumors subsequently annotated by a different dermatopathologist as either WD (grey), moderately-differentiated (MD (magenta)) or PD (teal). (B) Representative histology and FWE immunofluorescence of WD, MD and PD regions within cSCC originally classified as WD or PD (scale bars 1mm, 50µm (inset)). (C) Quantification of percent FWE positivity and FWE mean fluorescence intensity (MFI) in keratinocyte regions stratified by differentiation status (one way ANOVA with Tukey’s multiple comparisons test, p=*<0.05, **<0.01, ***<0.001, ****<0.0001). Annotations include adjacent normal skin (NS) and actinic keratosis (AK), in addition to WD, MD and PD cSCC.

## DISCUSSION

Flower (FWE) encodes a small, highly conserved canonical isoform, hFWE4, that contributes to polarized lamellar body secretion during normal epidermal barrier maturation (12). Here we show that, cSCC, which often retain tissue architecture resembling normal epidermis, also express FWE in highly differentiated cell populations. Genetic KO of *FWE* produces a histologic phenotype characterized by diminished keratohyalin-granule containing cells often accompanied by solid parakeratosis in keratin pearls. Transcriptomic analysis of these tumors suggests that this phenotype is most likely driven by abnormalities during end-stage cornification events including lamellar body biogenesis and secretion. Further indicating a functional role for FWE in cSCC differentiation, we also find that ectopic expression of the canonical hFWE4 isoform induces cell cycle arrest and elicits exit from the proliferative basal compartment. These findings are largely in line with our recent observations in normal keratinocytes and importantly underscore the critical, and previously unappreciated role of FWE in executing the final stages of terminal differentiation in the epidermis and its derivative tumors.

We find that despite a clear effect on keratinization patterns, knockout of *FWE* does not lead to significant changes in tumor growth or proliferation. This is likely owing to the fact that expression of *FWE* is largely restricted to cells in the upper suprabasal cell layers, while minimal expression is observed in the basal proliferative population that contributes most substantially to tumor growth. Indeed, *hFWE* KO does not appear to impact the proliferative potential of basal keratinocytes in cSCC. Intriguingly, despite the lack of effect on tumor volume, *hFWE* KO tumors exhibited significant increase in final tumor mass. Transcriptomic analysis suggests that there are likely significant alterations to the composition of cornified keratin pearls in the absence of *hFWE,* as genetic KO leads to marked dysregulation of genes involved in cornification including many involved in lamellar body function. We hypothesize that this compositional change in cornified layers is responsible for the mass discrepancy between genotypes, however more work is required to fully test this hypothesis.

Despite the lack of a proliferative effect when depleting endogenous *hFWE*, ectopic expression of hFWE4 elicits G1 arrest in culture and forces differentiation of cSCC cells in xenografts. We previously observed this same phenomenon in normal keratinocytes, where we found that chelating intracellular Ca^2+^ attenuated the hFWE4-dependent G1 arrest. It seems likely that this phenotype in cSCC cells is also dependent upon hFWE4-mediated liberation of intracellular Ca^2+^ stores, however more work is required to test this hypothesis. This phenomenon appears unique to keratinocytes as hFWE4 expression in breast and colon cancer cells provides a selective advantage (26). Differences in Ca^2+^ sensitivity between these cell types are the most obvious explanation for this discrepancy, however more work is needed to test this hypothesis. Regardless, the fact that ectopic expression of hFWE4 drives exit from the basal compartment raises the question of whether elevation of endogenous *hFWE* expression can contribute to the initiation of differentiation. Given that we are largely unable to detect substantial levels of FWE in the basal keratinocyte population in normal epidermis, it seems unlikely that this would occur during homeostatic regeneration of the epidermis, However, UV-irradiation or other exogenous stimuli could drive transient *hFWE* expression in small numbers of basal keratinocytes to elicit this response. In line with this idea, STAT3, a transcription factor implicated in the keratinocyte UVB response (27, 28), was recently found to bind to the *hFWE* promoter (29). Much work is required to test this hypothesis and importantly, more sensitive reporters or affinity reagents are required to capture these infrequent, transient elevations in expression if they occur.

Our analysis of FWE expression in human cSCC with highly variable keratinization patterns demonstrated that FWE typically labels the last several viable layers of keratinocytes preceding pearl formation regardless of pearl structure. We did however observe several tumors with well differentiated regions containing solid parakeratosis that exhibited notable absence of FWE signal in the cell layers directly adjacent to the keratinized pearl. This is in line with our observations that *hFWE* KO cSCCs more frequently produce this keratinization pattern that terminates with solid parakeratosis at the center of the keratin pearl. Whether these subtle alterations to keratinization pattern in human cSCCs are attributable to differences in *hFWE* expression is not yet clear and requires additional investigation.

Finally, given the conservation of its cornification-related function from normal epidermis to cSCC, we propose that immunostaining for FWE as a late differentiation marker may be a valuable prognostic tool in cSCC. While differentiation grading is often used to stratify risk associated with cSCC, grading criteria rely largely on appreciation of differentiation-related structures such as keratin pearls on routine histology. This produces considerable levels of both inter- and intra-observer variability, a problem highlighted in the present work by the fact that tumors initially identified as ‘poorly-differentiated’ were largely composed of ‘moderately-differentiated’ keratinocyte regions. Analysis of patient tumors suggests that both FWE positive area and FWE intensity reliably distinguish ‘well-differentiated’ from ‘moderately-‘ or ‘poorly-differentiated’ cSCC cells. While there is a trend towards reduction in both of these metrics from ‘moderately-’ to ‘poorly-differentiated’ regions as well, this finding was not significant. Additional work using larger sample cohorts are needed to determine whether FWE signal could be used to distinguish between ‘moderately-‘ and ‘poorly-differentiated’ tumors. While we apply immunofluorescence in the present study, future work may seek to develop a reliable immunohistochemical assay for FWE to more easily integrate into standard pathology workflows.

## METHODS

### Cell Culture

Cutaneous squamous cell carcinoma cell lines, SCC-13 (18, 19), SCC-12b.2 (18, 19) and COLO 16 (20) were grown in complete high-glucose DMEM (Gibco 11965118) supplemented with 10%FBS and 1% penicillin-streptomycin, or EpiLife supplemented with HKGS (Gibco S0015), 1% penicillin-streptomycin and either 0.4mM or 1.4mM CaCl_2_ where indicated. Immortalized N/TERT-2G keratinocytes were grown in EpiLife media supplemented as described. All cells were maintained in a humidified incubator at 37°C, 5% CO_2_.

For induction of terminal differentiation in cell culture experiments, cells were seeded at 100% confluency, switched to 1.4mM CaCl_2_ after 24h and refed daily until the indicated timepoint. Cell culture lysates were prepared in RIPA buffer supplemented with 1x HALT Protease and Phosphatase Inhibitor (Thermo Fisher 78425). Following protein estimation, equivalent protein mass was denatured in 4x Laemmli buffer and separated on an SDS-PAGE gel before transfer to nitrocellulose membrane for immunoblotting using standard techniques. Primary antibodies used for immunoblotting included anti-α/β Tubulin (Cell Signaling Technology 2148), anti-Loricrin (Biolegend Poly19051) and anti-FWE (Cell Signaling Technology 28284) or anti-FWE (16).

### Molecular Cloning, Lentivirus and Stable Cell Line Generation

pLenti CMV-MCS-EGFP-SV40-Puro (Addgene #73582) was XbaI/BamHI digested and the purified backbone was used as the base vector for generating lentiviral hFWE4 expression constructs. Briefly, two-fragment In-Fusion Snap Assembly (Takara Bio 638947) was performed to insert a PCR amplified EGFP cassette in-frame with P2A-hFWE4-3XFLAG synthesized as a double-stranded gBlocks (IDT). Plasmidsaurus whole-plasmid sequencing was used to validate final vector sequence.

Lentivirus was packaged as previously described (17). Briefly, HEK293FT (Invitrogen) were co-transfected with the transfer vector of interest along with psPAX2 and pMD2.G packaging vectors following standard techniques. Supernatants were concentrated using Lenti-X concentrator (Takara Bio 631231). Indicated cells were ‘spinfected’ in multi-well plates by combining cells in suspension with lentiviral particles and 10 μg/ml polybrene and centrifuging at 1,000 x g for 1h at 37°C. Functional lentiviral titers were quantified via flow cytometry using this same ‘spinfection’ technique to allow equivalent transduction efficiency across constructs in experiments.

For stable expression of lentiviral constructs in SCC-13 cells used for xenografting and long-term cell culture experiments, FACS was used to sort cells to purity following lentiviral infection. Cells were subsequently maintained in media containing 1 μg/ml puromycin to maintain selective pressure.

### Cell Cycle Profiling

SCC-13, SCC-12b.2 or COLO 16 cells were ‘spinfected’ with EGFP or EGFP 2A hFWE4 encoding lentivirus as described above. Cells were subcultured 48h after spinfection and subsequently pulsed either 24h (COLO 16) or 48h (SCC-13, SCC-12b.2) after seeding with 10mM EdU for 1h. Cells were collected, fixed and processed for Click-It EdU labeling (Invitrogen C10419) and FxCycle Violet (Invitrogen R37166) DNA content labeling according to manufacturers instructions. Cell cycle data was acquired on a YETI flow cytometer with EGFP+ singlets analyzed for EdU vs FxCycle signal using Everest (BioRad) software.

### CRISPR/Cas9 Editing and Clonal Cell Line Generation

*hFWE* KO and WT clonal cell lines were derived through limiting dilution cloning of edited cell pools generated by Synthego through electroporation of Cas9:sgRNA RNP targeting Exon 2 (5’UGAACAUCGCGGCCGGCGUG3’) or Cas9 only. On expansion, gDNA was isolated (Qiagen QiaAMP DNA Blood Mini 51104) and the edited site was Sanger sequenced (Genewiz). The Interference of CRISPR Edits (ICE) tool (Synthego) was used to analyze sequencing traces for clonality. Sequencing primers can be found in Supplementary Table 2.

### Subcutaneous Xenografts

For subcutaneous xenografts of *hFWE* WT or KO cells, 1*10^6^ SCC13 cells were resuspended in 100µl of cold 1:1 sterile PBS:Matrigel (Corning 356237) and subsequently injected into the subcutaneous space on the dorsal flank of a NOD-Prkdc^em26Cd52^Il2rg^em26Cd22^/NjuCrl (NCG) mouse (Charles River 572). For subcutaneous xenografts of 1:1 cocultures, 0.5*10^6^ SCC13 cells stably expressing EGFP alone or EGFP P2A hFWE4 were mixed with 0.5*10^6^ SCC13 cells stably expressing mCherry. An aliquot of the coculture mixture was subject to flow cytometry to determine the starting ratio of GFP+:mCherry+ cells. The final cocultures containing 1*10^6^ total cells were then resuspended in 1:1 sterile PBS:Matrigel and inoculated into NCG mice as described above. Tumor volumes were measured with electronic calipers twice weekly and animal weights were monitored at regular intervals. At necropsy, tumors were weighed before partitioning for RNA isolation or histological processing. RNA was isolated from flash frozen samples using Qiagen RNeasy Mini Kit (Qiagen 74104).

### RNA Sequencing

For preparation of sequencing libraries, 500ng of RNA isolated from flash frozen tumors was used with the NEBNext Ultra II Directional RNA Library Prep kit for Illumina along with NEBNext Poly(A) mRNA Magnetic Isolation Module according to the manufacturers instructions. Library quality and quantity were assessed using RNA ScreenTape Assay (Agilent) and Agilent 4200 TapeStation System following the manufacturers protocol. 3 nM equimolar pooled libraries were sequence on the Illumina NextSeq 2000 system (Illumina) using NextSeq 1000/2000 P2 Reagent Cartridge 100 cycles and P2 Flow Cell (Illumina).

Analysis of bulk RNA-seq data was performed in PartekFlow. Adapter-trimmed reads were aligned to hg38 using STAR (30) and quantified to ‘hg38-Ensembl Transcripts release 105’ using PartekE/M. Gene counts were normalized by median ratio, and normalized counts were used for differential expression analysis using DESeq2 (31).

ShinyGO 0.82 (32) was used to perform GO:BP and GO:CC analysis on downregulated (FDR<0.05, l2fc>1) genes. Pathway size was restricted to a minimum of 100 genes for GO:BP and 10 genes for GO:CC, with maximum of 5000 genes for both.

### Histology, Immunofluorescence, Microscopy and Image Analysis

FFPE sections from deidentified human tumors or subcutaneous xenografts were subject to H&E or immunofluorescence staining using standard protocols. For immunofluorescence, rehydrated sections were subject to citrate-based antigen retrieval (VectorLabs) prior to blocking and primary antibody incubation in 1%BSA/5% goat serum/0.3% TritonX-100. Samples were labeled with fluorophore-conjugated secondary antibodies and nuclei were counterstained with Hoechst 33342 before mounting coverslips with VectaShield Antifade mounting medium (VectorLabs). Details regarding primary and secondary antibodies used in immunofluorescence assays can be found in Supplementary Table 3.

Imaging acquisition was performed on an Olympus VS120-S6-W slide scanning system equipped with DAPI, FITC, TRITC and Cy5 filter cubes. Quantitative analysis of Loricrin and Filaggrin immunofluorescence was performed using Olympus VS-Desktop software. Briefly, eight 60,000 µm^2^ regions were annotated per tumor and a common intensity threshold was applied for each marker based on no primary controls. Percent positive thresholded area within each annotated region was extracted and an average for each tumor was calculated.

For quantitative analysis of FWE immunofluorescence, digitized H&E scans were first annotated with regions of normal skin, AK, well-differentiated SCC, moderately-differentiated SCC and poorly-differentiated SCC by a dermatopathologist blinded to the immunofluorescence signal. Annotations were copied forward onto digitized immunofluorescence for analysis in QuPath-0.5.1-arm64 (33). Mean fluorescence intensity (MFI) values for each annotation were extracted. For calculation of percent positive area, a common threshold was applied to detect signal above background observed in basal keratinocytes of adjacent normal skin. Thresholded area was then extracted for each annotation.

## ACKNOWLEDGEMENTS

We acknowledge additional support provided by members of the Advanced Microscopy, Flow Cytometry, Innovative Genomics and Bioinformatics core facilities as well as the Integrated Biomedical Imaging Facility (IBIF) at Creighton University. This work was supported by the State of Nebraska LB595, LB506 and LB606.

## REFERENCES

1. Simpson, C. L., Patel, D. M., and Green, K. J. (2011) Deconstructing the skin: cytoarchitectural determinants of epidermal morphogenesis. Nat Rev Mol Cell Biol. 12, 565–580

2. Candi, E., Schmidt, R., and Melino, G. (2005) The cornified envelope: a model of cell death in the skin. Nat Rev Mol Cell Biol. 6, 328–340

3. Paller, A. S., Teng, J., Mazereeuw-Hautier, J., Hernández-Martín, Á., Tournier, C. G., Hovnanian, A., Aldwin-Easton, M., Tadini, G., Janice, S., Sprecher, E., Malovitski, K., Ishida-Yamamoto, A., Choate, K., Akiyama, M., O’Toole, E. A., Fischer, J., Bodemer, C., Gostynski, A., and Schmuth, M. (2025) Syndromic epidermal differentiation disorders: New classification towards pathogenesis-based therapy. Br J Dermatol. 10.1093/bjd/ljaf123

4. Akiyama, M., Choate, K., Hernandez-Martin, A., Aldwin-Easton, M., Bodemer, C., Gostyński, A., Hovnanian, A., Ishida-Yamamoto, A., Malovitski, K., O’Toole, E. A., Paller, A. S., Schmuth, M., Schwartz, J., Sprecher, E., Teng, J. M. C., Granier Tournier, C., Mazereeuw-Hautier, J., Tadini, G., and Fischer, J. (2025) Nonsyndromic epidermal differentiation disorders: New classification and nomenclature based on disease-associated genes leading to targeted therapy. Br J Dermatol. 10.1093/bjd/ljaf154

5. Lopez-Pajares, V., Yan, K., Zarnegar, B. J., Jameson, K. L., and Khavari, P. A. (2013) Genetic pathways in disorders of epidermal differentiation. Trends in Genetics. 29, 31–40

6. Sánchez-Danés, A., and Blanpain, C. (2018) Deciphering the cells of origin of squamous cell carcinomas. Nat Rev Cancer. 18, 549–561

7. Que, S. K. T., Zwald, F. O., and Schmults, C. D. (2018) Cutaneous squamous cell carcinoma: Incidence, risk factors, diagnosis, and staging. Journal of the American Academy of Dermatology. 78, 237–247

8. Thompson, A. K., Kelley, B. F., Prokop, L. J., Murad, M. H., and Baum, C. L. (2016) Risk Factors for Cutaneous Squamous Cell Carcinoma Recurrence, Metastasis, and Disease-Specific Death: A Systematic Review and Meta-analysis. JAMA Dermatol. 152, 419–428

9. Prezzano, J. C., Scott, G. A., Lambert Smith, F., Mannava, K. A., and Ibrahim, S. F. (2021) Concordance of Squamous Cell Carcinoma Histologic Grading Among Dermatopathologists and Mohs Surgeons. Dermatologic Surgery. 47, 1433

10. Nash, J., Shahwan, K. T., Chung, C., Abidi, N., Gokun, Y., Pan, X., and Carr, D. R. (2022) Grading of differentiation in cutaneous squamous cell carcinoma: Evaluation of interrater and intrarater reliability. Journal of the American Academy of Dermatology. 87, 895–897

11. Diede, C., Walker, T., Carr, D. R., and Shahwan, K. T. (2024) Grading differentiation in cutaneous squamous cell carcinoma: a review of the literature. Arch Dermatol Res. 316, 434

12. Rudd, J. C., Smits, J. P. H., Kuwong, P. T., Johnson, R. E., Monga, L. M. N., van Vlijmen-Willems, I. M. J. J., Porter, G. L., Halloran, P. O., Sharma, K., Schmidt, K. N., Kumar, V., Madson, J. G., Sarkar, M. K., van den Bogaard, E. H., Grunkemeyer, J. A., Gudjonsson, J. E., Wong, S. Y., Simpson, C. L., and Hansen, L. A. (2025) Flower dependent trafficking of lamellar bodies facilitates maturation of the epidermal barrier. Nat Commun. 16, 6892

13. Yao, C.-K., Lin, Y. Q., Ly, C. V., Ohyama, T., Haueter, C. M., Moiseenkova-Bell, V. Y., Wensel, T. G., and Bellen, H. J. (2009) A Synaptic Vesicle-Associated Ca2+ Channel Promotes Endocytosis and Couples Exocytosis to Endocytosis. Cell. 138, 947–960

14. Yao, C.-K., Liu, Y.-T., Lee, I.-C., Wang, Y.-T., and Wu, P.-Y. (2017) A Ca2+ channel differentially regulates Clathrin-mediated and activity-dependent bulk endocytosis. PLoS Biol. 15, e2000931

15. Li, T.-N., Chen, Y.-J., Lu, T.-Y., Wang, Y.-T., Lin, H.-C., and Yao, C.-K. (2020) A positive feedback loop between Flower and PI(4,5)P2 at periactive zones controls bulk endocytosis in Drosophila. Elife. 9, e60125

16. Chang, H.-F., Mannebach, S., Beck, A., Ravichandran, K., Krause, E., Frohnweiler, K., Fecher-Trost, C., Schirra, C., Pattu, V., Flockerzi, V., and Rettig, J. (2018) Cytotoxic granule endocytosis depends on the Flower protein. J Cell Biol. 217, 667–683

17. Rudd, J. C., Maity, S., Grunkemeyer, J. A., Snyder, J. C., Lovas, S., and Hansen, L. A. (2023) Membrane structure and internalization dynamics of human Flower isoforms hFWE3 and hFWE4 indicate a conserved endocytic role for hFWE4. Journal of Biological Chemistry. 10.1016/j.jbc.2023.104945

18. Rheinwald, J. G., and Beckett, M. A. (1980) Defective terminal differentiation in culture as a consistent and selectable character of malignant human keratinocytes. Cell. 22, 629–632

19. Rheinwald, J. G., and Beckett, M. A. (1981) Tumorigenic keratinocyte lines requiring anchorage and fibroblast support cultured from human squamous cell carcinomas. Cancer Res. 41, 1657–1663

20. Moore, G. E., Merrick, S. B., Woods, L. K., and Arabasz, N. M. (1975) A Human Squamous Cell Carcinoma Cell Line1. Cancer Res. 35, 2684–2688

21. Shin, J.-M., Chang, I.-K., Lee, Y.-H., Yeo, M.-K., Kim, J.-M., Sohn, K.-C., Im, M., Seo, Y.-J., Kim, C.-D., Lee, J.-H., and Lee, Y. (2016) Potential Role of S100A8 in Cutaneous Squamous Cell Carcinoma Differentiation. Ann Dermatol. 28, 179–185

22. Alonso-Lecue, P., de Pedro, I., Coulon, V., Molinuevo, R., Lorz, C., Segrelles, C., Ceballos, L., López-Aventín, D., García-Valtuille, A., Bernal, J. M., Mazorra, F., Pujol, R. M., Paramio, J., Ramón Sanz, J., Freije, A., Toll, A., and Gandarillas, A. (2017) Inefficient differentiation response to cell cycle stress leads to genomic instability and malignant progression of squamous carcinoma cells. Cell Death Dis. 8, e2901–e2901

23. Holmes, T. R., Al Matouq, J., Holmes, M., Sioda, N., Rudd, J. C., Bloom, C., Nicola, L., Palermo, N. Y., Madson, J. G., Lovas, S., and Hansen, L. A. (2021) Targeting 14-3-3ε activates apoptotic signaling to prevent cutaneous squamous cell carcinoma. Carcinogenesis. 42, 232–242

24. Holmes, T. R., Al-Matouq, J., Holmes, M., Nicola, L., Rudd, J. C., Lovas, S., and Hansen, L. A. (2020) Targeting 14-3-3ε-CDC25A interactions to trigger apoptotic cell death in skin cancer. Oncotarget. 11, 3267–3278

25. Rubin, A. L., Parenteau, N. L., and Rice, R. H. (1989) Coordination of keratinocyte programming in human SCC-13 squamous carcinoma and normal epidermal cells. Journal of Cellular Physiology. 138, 208–214

26. Madan, E., Pelham, C. J., Nagane, M., Parker, T. M., Canas-Marques, R., Fazio, K., Shaik, K., Yuan, Y., Henriques, V., Galzerano, A., Yamashita, T., Pinto, M. A. F., Palma, A. M., Camacho, D., Vieira, A., Soldini, D., Nakshatri, H., Post, S. R., Rhiner, C., Yamashita, H., Accardi, D., Hansen, L. A., Carvalho, C., Beltran, A. L., Kuppusamy, P., Gogna, R., and Moreno, E. (2019) Flower isoforms promote competitive growth in cancer. Nature. 572, 260–264

27. Bito, T., Sumita, N., Masaki, T., Shirakawa, T., Ueda, M., Yoshiki, R., Tokura, Y., and Nishigori, C. (2010) Ultraviolet light induces Stat3 activation in human keratinocytes and fibroblasts through reactive oxygen species and DNA damage. Exp Dermatol. 19, 654–660

28. Sano, S., Chan, K. S., Kira, M., Kataoka, K., Takagi, S., Tarutani, M., Itami, S., Kiguchi, K., Yokoi, M., Sugasawa, K., Mori, T., Hanaoka, F., Takeda, J., and DiGiovanni, J. (2005) Signal transducer and activator of transcription 3 is a key regulator of keratinocyte survival and proliferation following UV irradiation. Cancer Res. 65, 5720–5729

29. Madan, E., Palma, A. M., Vudatha, V., Kumar, A., Bhoopathi, P., Wilhelm, J., Bernas, T., Martin, P. C., Bilolikar, G., Gogna, A., Peixoto, M. L., Dreier, I., Araujo, T. F., Garre, E., Gustafsson, A., Dorayappan, K. D. P., Mamidi, N., Sun, Z., Yekelchyk, M., Accardi, D., Olsen, A. L., Lin, L., Titelman, A. A., Bianchi, M., Jessmon, P., Farid, E. A., Pradhan, A. K., Neufeld, L., Yeini, E., Maji, S., Pelham, C. J., Kim, H., Oh, D., Rolfsnes, H. O., Marques, R. C., Lu, A., Nagane, M., Chaudhary, S., Gupta, K., Gogna, K. C., Bigio, A., Bhoopathi, K., Mannangatti, P., Achary, K. G., Akhtar, J., Belião, S., Das, S., Correia, I., da Silva, C. L., Fialho, A. M., Poellmann, M. J., Javius-Jones, K., Hawkridge, A. M., Pal, S., Shree, K. S., Rakha, E. A., Khurana, S., Xiao, G., Zhang, D., Rijal, A., Lyons, C., Grossman, S. R., Turner, D. P., Pillappa, R., Prakash, K., Gupta, G., Robinson, G. L. W. G., Koblinski, J., Wang, H., Singh, G., Singh, S., Rayamajhi, S., Bacolod, M. D., Richards, H., Sayeed, S., Klein, K. P., Chelmow, D., Satchi-Fainaro, R., Selvendiran, K., Connolly, D., Thorsen, F. A., Bjerkvig, R., Nephew, K. P., Idowu, M. O., Kühnel, M. P., Moskaluk, C., Hong, S., Redmond, W. L., Landberg, G., Lopez-Beltran, A., Poklepovic, A. S., Sanyal, A., Fisher, P. B., Church, G. M., Menon, U., Drapkin, R., Godwin, A. K., Luo, Y., Ackermann, M., Tzankov, A., Mertz, K. D., Jonigk, D., Tsung, A., Sidransky, D., Trevino, J., Saavedra, A. P., Winn, R., Won, K. J., Moreno, E., and Gogna, R. (2024) Ovarian tumor cells gain competitive advantage by actively reducing the cellular fitness of microenvironment cells. Nat Biotechnol. 10.1038/s41587-024-02453-3

30. Dobin, A., Davis, C. A., Schlesinger, F., Drenkow, J., Zaleski, C., Jha, S., Batut, P., Chaisson, M., and Gingeras, T. R. (2013) STAR: ultrafast universal RNA-seq aligner. Bioinformatics. 29, 15–21

31. Love, M. I., Huber, W., and Anders, S. (2014) Moderated estimation of fold change and dispersion for RNA-seq data with DESeq2. Genome Biology. 15, 550

32. Ge, S. X., Jung, D., and Yao, R. (2020) ShinyGO: a graphical gene-set enrichment tool for animals and plants. Bioinformatics. 36, 2628–2629

33. Bankhead, P., Loughrey, M. B., Fernández, J. A., Dombrowski, Y., McArt, D. G., Dunne, P. D., McQuaid, S., Gray, R. T., Murray, L. J., Coleman, H. G., James, J. A., Salto-Tellez, M., and Hamilton, P. W. (2017) QuPath: Open source software for digital pathology image analysis. Sci Rep. 7, 16878

