## Supplementary Figures for "The human Flower isoform hFWE4 facilitates cornification in cutaneous squamous cell carcinoma"

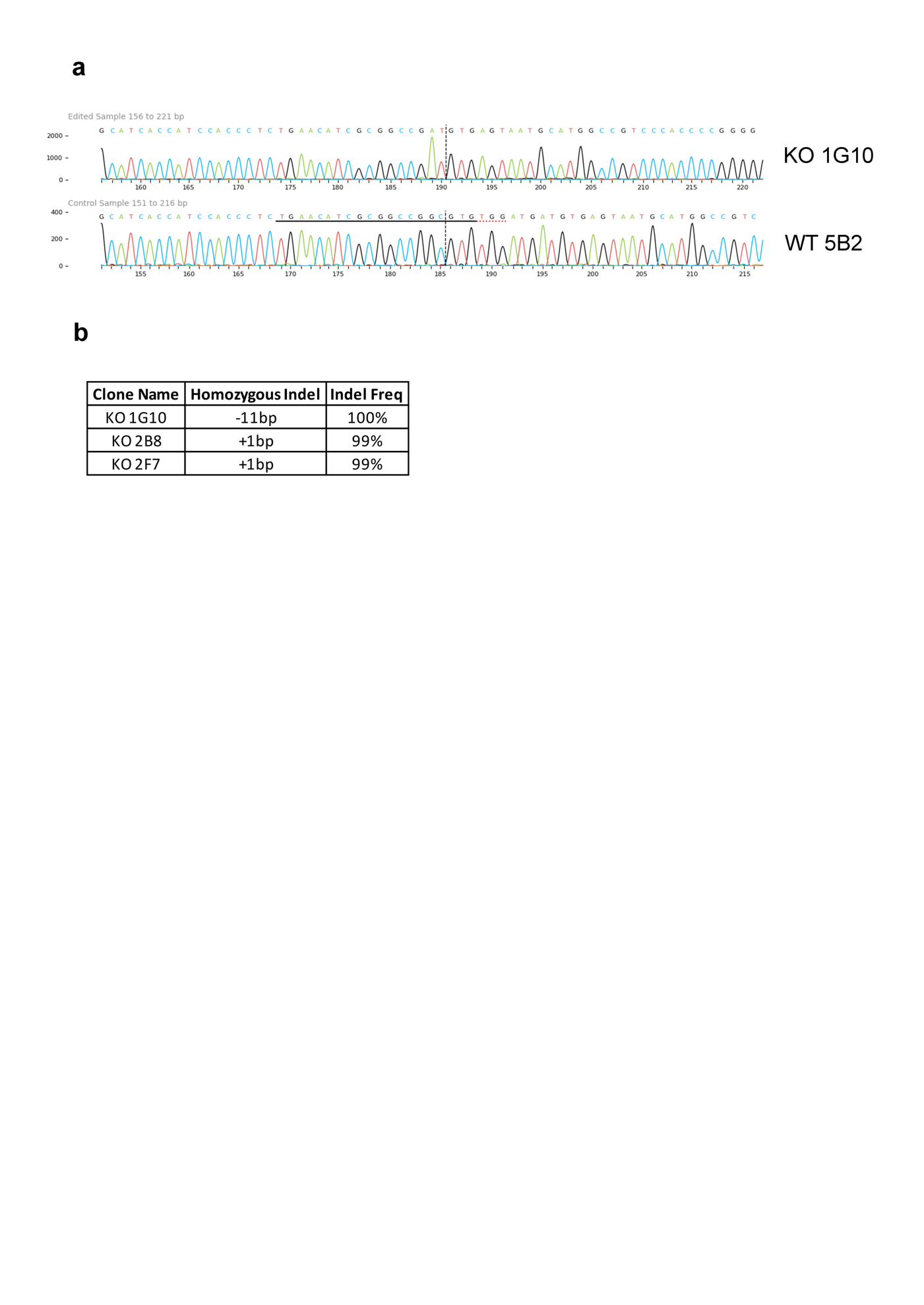
**Supplemental Figure 1. Sanger sequencing validation of *hFWE* KO in SCC-13 cells. (a)** Aligned sanger sequencing traces from interference of CRISPR edits (ICE) analysis. KO clone 1G10 is aligned against WT 5B2 for representation. **(b)** Predicted mutation frequency of all KO clones used in xenograft experiments.

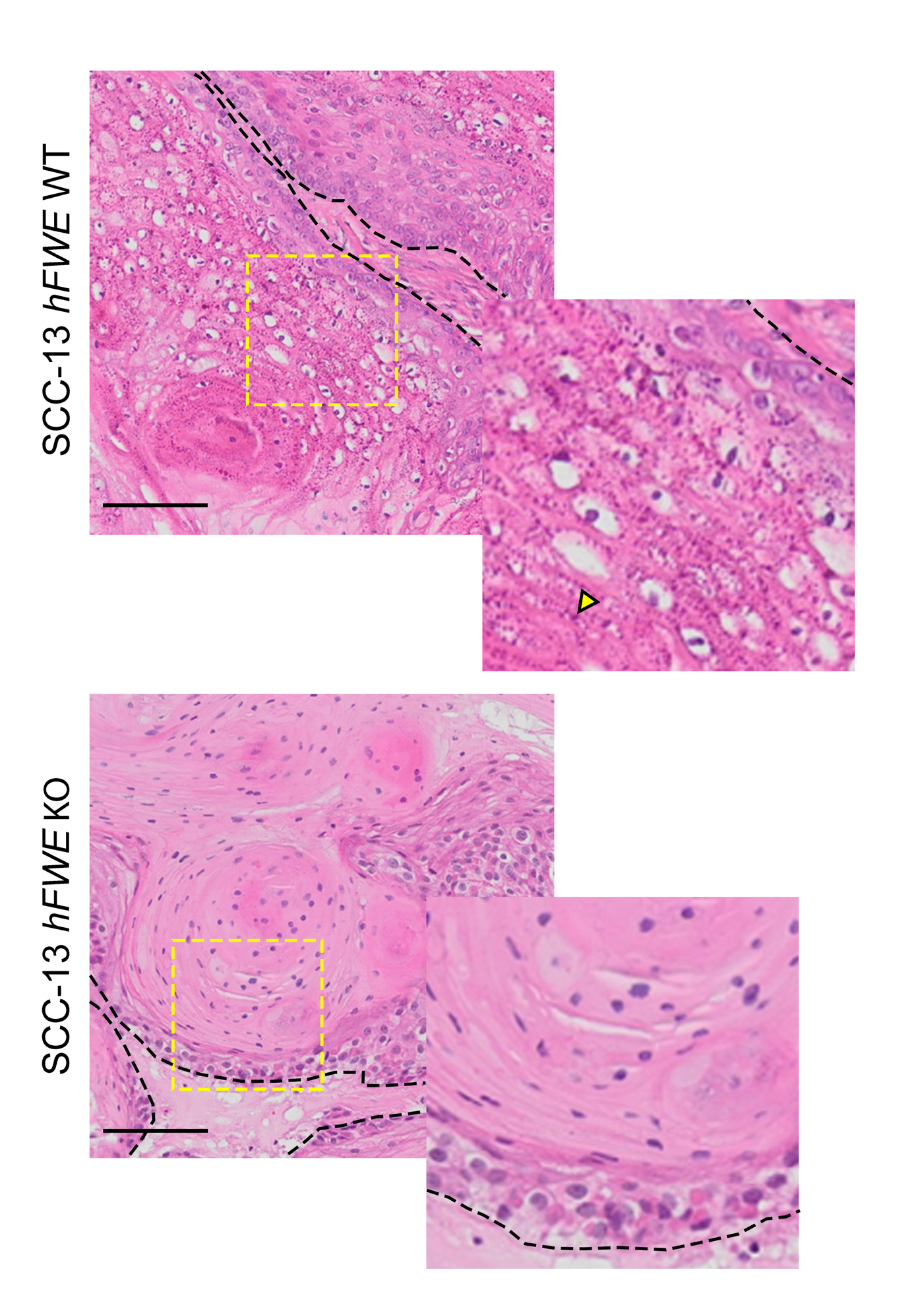
**Supplemental Figure 2. Granular and cornified layer changes in *hFWE* KO SCC-13 xenografts.** Higher magnification images from Fig 2E showing representative alterations in granular and cornified layers of *hFWE* KO SCC-13 xenografts (scale bar 100µm). Dashed yellow box indicates inset area, dashed black line indicates basement membrane. Note the absence of keratohyalin granules (yellow arrowhead) and presence of solid parakeratosis in KO tumors.

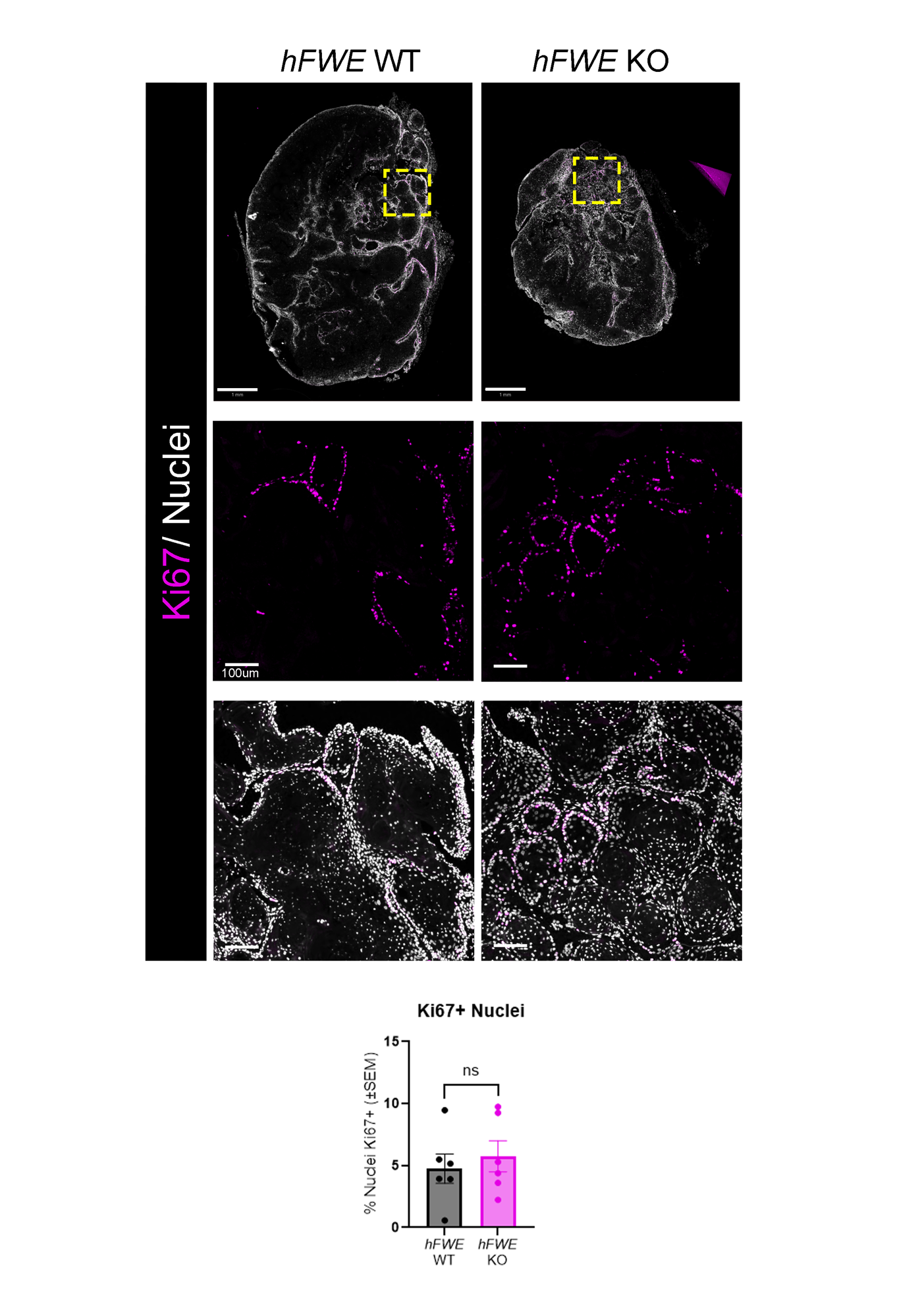
**Supplemental Figure 3. *hFWE* KO does not affect proliferation in SCC-13 xenografts**. (**a**) Representative immunofluorescence for Ki67 in *hFWE* WT and KO SCC-13 xenografts (scale bars 1mm, 100um (inset)). (**b**) Quantification of percentage of nuclear area positive for Ki67 across tumor genotype (p>0.05, two-tailed independent t-test).

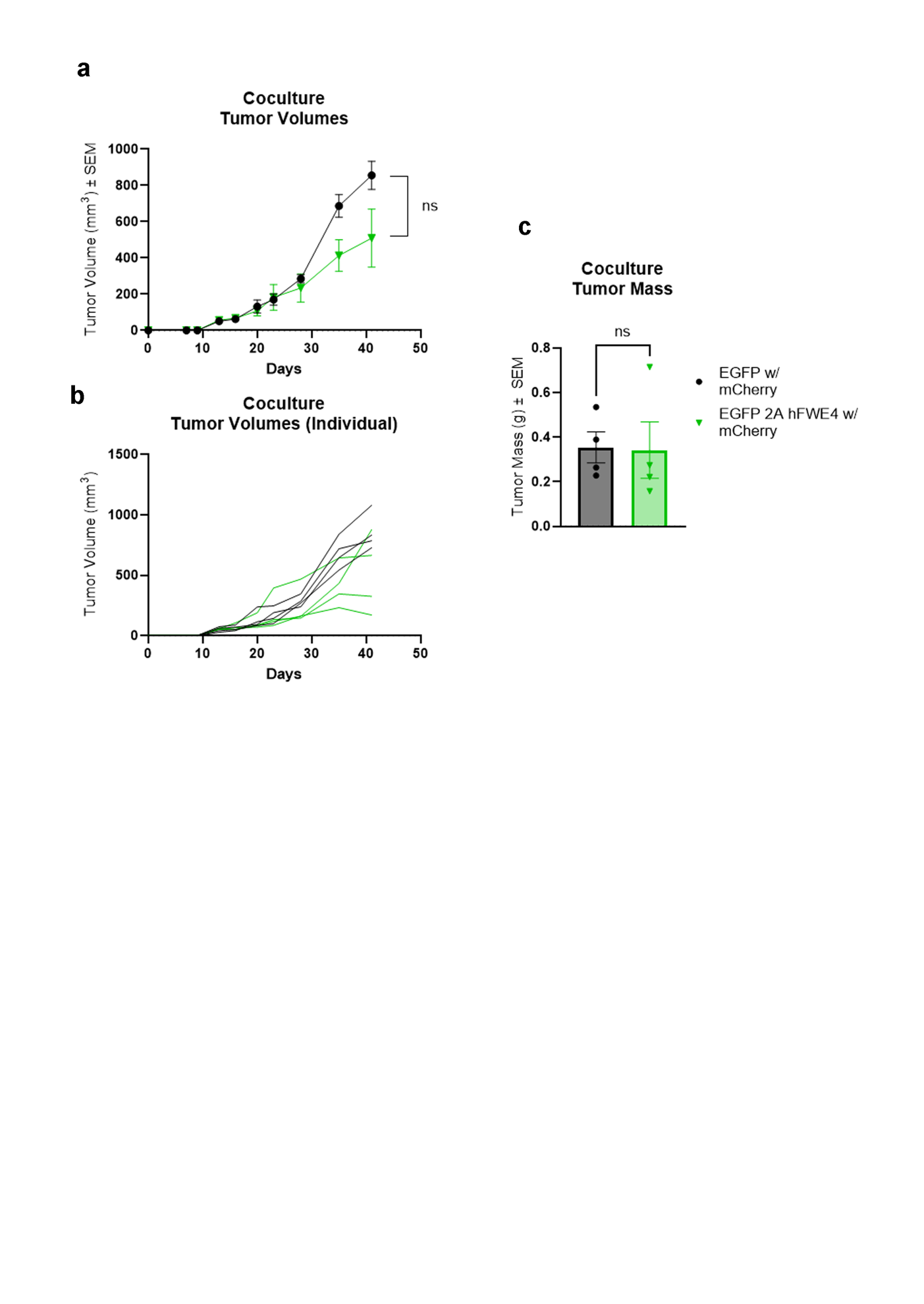
**Supplemental Figure 4. Mixed hFWE4 xenografts do not exhibit reduction in tumor volume or mass.** Quantification of mean (**a**) or individual (**b**) tumor volumes from mixed SCC-13 xenografts (p<0.05, multiple unpaired t-tests with post-hoc Holm-Sidak correction). (**c**) Quantification of final tumor mass from mixed SCC-13 xenografts (p<0.05, two-tailed independent t-test).
